# Single-Cell Analysis Reveals Cryptic Prophage Protease LfgB Protects *Escherichia coli* During Oxidative Stress by Cleaving Antitoxin MqsA

**DOI:** 10.1101/2023.09.23.559088

**Authors:** Laura Fernández-García, Xinyu Gao, Michael E. Battisti, Joy Kirigo, Rodolfo García-Contreras, Maria Tomas, Yunxue Guo, Xiaoxue Wang, Thomas K. Wood

**Affiliations:** Department of Chemical Engineering, Pennsylvania State University, University Park, Pennsylvania, 16802-4400, USA; Microbiology Translational and Multidisciplinary (MicroTM)-Research Institute Biomedical A Coruña (INIBIC) and Microbiology Department of Hospital A Coruña (CHUAC), University of A Coruña (UDC); Key Laboratory of Tropical Marine Bio-resources and Ecology, Guangdong Key Laboratory of Marine Materia Medica, Innovation Academy of South China Sea Ecology and Environmental Engineering, South China Sea Institute of Oceanology, Chinese Academy of Sciences, No.1119, Haibin Road, Nansha District, Guangzhou 511458, China; University of Chinese Academy of Sciences, Beijing 100049, China; Departamento de Microbiología y Parasitología, Facultad de Medicina, Universidad Nacional Autónoma de México, Mexico City, Mexico; Southern Marine Science and Engineering Guangdong Laboratory (Guangzhou), No.1119, Haibin Road, Nansha District, Guangzhou 511458, China

## Abstract

Although toxin/antitoxin (TA) systems are ubiquitous, beyond phage inhibition and mobile element stabilization, their role in host metabolism is obscure. One of the best-characterized TA systems is MqsR/MqsA of *Escherichia coli*, which has been linked previously to protecting this gastrointestinal species during the stress it encounters from the bile salt deoxycholate as it colonizes humans. However, some recent whole-population studies have challenged the role of toxins such as MqsR in bacterial physiology, since the *mqsRA* locus is induced over a hundred-fold during stress, but a phenotype was not found upon its deletion. Here, we investigate further the role MqsR/MqsA by utilizing single cells and demonstrate that upon oxidative stress, the TA system MqsR/MqsA has a heterogeneous effect on the transcriptome of single cells. Furthermore, we discovered that MqsR activation leads to induction of the poorly-characterized *yfjXY ypjJ yfjZF* operon of cryptic prophage CP4-57. Moreover, deletion of *yfjY* makes the cells sensitive to H_2_O_2_, acid, and heat stress, and this phenotype was complemented. Hence, we recommend *yfjY* be renamed to *lfgB* (less fatality <u>g</u>ene <u>B</u>). Critically, MqsA represses *lfgB* by binding the operon promoter, and LfgB is a protease that degrades MqsA to derepress *rpoS* and facilitate the stress response. Therefore, the MqsR/MqsA TA system facilitates the stress response through cryptic phage protease LfgB.

## INTRODUCTION

Toxin/antitoxin systems are encoded in the genomes of nearly all archaea and bacteria and are classified as eight main types based on how the antitoxin inactivates the toxin^55^. We discovered that phage inhibition is one of the primary physiological roles of TA systems and determined the mechanism is toxin induction via host transcription shutdown by the attacking phage^37^; these results were confirmed 25 years later^18^. TA systems also stabilize mobile genetic elements^16, 35, 46, 58^. Beyond these two functions, there is controversy regarding the physiological roles of TA systems^44^.

The MqsR/MqsA TA system was discovered as induced in a biofilm transcriptome study^39^ and shown to be a TA system using the structures of the toxin (MqsR) and antitoxin (MqsA)^5^. MqsR degrades mRNA with the 5’-GCU site^60^, and MqsA was found to regulate not only its own promoter but to repress the oxidative stress response via DNA binding at a palindrome upstream of the stress response sigma factor RpoS^51^ and to repress curli synthesis by binding to the promoter of the gene that encodes the master biofilm regulator CsgD^45^. Moreover, MqsR/MqsA controls the TA system GhoT/GhoS as a cascade^54^ and helps *Escherichia coli* colonize the gastrointestinal tract by surviving bile acid stress^27^; activation of toxin MqsR during bile stress leads to degradation of YgiS mRNA, and this transcript encodes a periplasmic protein that promotes bile uptake. Furthermore, several groups have linked MqsR/MqsA to antibiotic tolerance based upon deletion of *mqsR*^24, 32, 59^, and MqsR/MqsA have been linked to heat shock^40^, biofilm formation^43^, nitrogen starvation^12^, and nitric oxide^36^ in *E. coli*, and copper stress^34^, vesicles^42^, and biofilm formation^29^ in *Xylella fastidiosa* as well as biofilm formation in *Pseudomonas fluorescens* ^56^ and persistence and biofilm formation in *P. putida*^47^.

In contrast to these myriad results with MqsR/MqsA, a report based on negative results claimed that the *E. coli* MqsR/MqsA TA system has no role in stress resistance, based on a lack of induction of the *mqsRA* locus and a lack of phenotype upon deleting *mqsRA*^13^. Strikingly, these transcription results were invalidated within a few months as *mqsRA* transcription in the wild-type strain was shown to increase by over 181-fold during amino acid stress and 90-fold during oxidative stress^30^. This work^30^ also claimed there was no physiological effect of deleting *mqsRA*, but, unfortunately, they utilized a TA deletion strain that has substantial non-related mutations, including large chromosomal inversions^17^; utilization of TA deletion strains with many coding errors beyond those of the TA systems has led to notorious retractions in the TA field, as we have summarized previously^57^. Critically, their clam^30^ of a lack of a physiological role of MqsR/MqsA was undercut by their later results which showed MqsR/MqsA/MqsC inhibited T2 phage^48^. We have confirmed these results and shown that phage inhibition by MqsRAC induces persistence rather than abortive infection^11^. Furthermore, these groups^13, 30^ used strains with *both* MqsR and MqsA inactivated rather than studying the effect of either the toxin or antitoxin alone; i.e., MqsR/MqsA work *together* during the oxidative stress response.

Based on these inconsistencies, we hypothesized that a better approach, due to heterogeneous gene expression^26^, would be to investigate the impact of MqsR/MqsA on cell physiology by monitoring the transcriptome of *single cells* since all previous studies have been based on population averages. Single cell transcriptomic studies have been initiated by several labs^4, 10, 20, 26, 33^, and here, we utilized the high-throughput microfluidic approach that relies on labeling each transcript with unique 50 nt single strand DNA probes to determine the impact of inactivation of MqsR/MqsA during oxidative stress^33^. We chose oxidative stress as the representative insult to cells since both anaerobes and aerobes must deal with this nearly-universal stress^22^, and MqsA has been shown to negatively regulate the oxidative stress response^51^. Using this approach, we determined that the *lfgABCDE* operon (formerly the uncharacterized operon *yfjXY ypjJ yfjZF*) of cryptic prophage CP4-57 is induced in single cells and that LfgB is a protease that is repressed by antitoxin MqsA and degrades MqsA to activate the *E. coli* stress response through sigma factor RpoS.

## RESULTS

### Antitoxin MqsA reduces the population stress response

We first investigated whether deleting an unmarked *mqsRA* mutation affected the response of *E. coli* to oxidative stress (20 mM H_2_O_2_ for 10 min) and found that, for the whole population, the wild-type cells were *more* sensitive to H_2_O_2_ (85 ± 15% death for the wild-type vs. 55 ± 10% for *mqsRA*). Similar population-wide results were seen with acid stress (pH 2.5 for 10 min for four cycles) where the wild-type strain was 64 times more sensitive. These results agree well with our previous results showing antitoxin MqsA represses *rpoS* by binding at a palindrome to help regulate stress resistance^51^. Moreover, our results suggest that inactivating toxin MqsR should reduce viability by elevating MqsA concentrations, since the additional antitoxin MqsA will repress *rpoS*, and as expected, when *mqsR* is deleted, cells are 14 ± 6 times more sensitive than the wild-type to oxidative stress. Therefore, the *mqsRA* mutant is better prepared to withstand oxidative and acid stress as its stress response via RpoS is activated due to the absence of the repressor MqsA.

### Single-cell analysis reveals LfgB increases cell viability during oxidative stress

Using single cells, we then investigated further the role of MqsR/MqsA during oxidative stress by comparing the wild-type strain vs. the unmarked *mqsRA* mutant in single cells. Utilizing 20 mM H_2_O_2_ for 10 min, we found (**Table 1**) that several cryptic prophage genes are induced in the wild-type strain relative to the *mqsRA* mutant, including *lfgA* of the *lfgABCDE* operon; previously LfgD (YfjZ) of this operon was shown by us to enhance MqsR toxicity^15^. Furthermore, the induction of two genes that encode heat-shock proteins (*ibpAB*) and one gene that encodes an osmotic stress response protein (*yciF*) served as positive controls for our single-cell analysis.

**Table 1.** Impact on gene expression after inactivating the MqsR/MqsA toxin/antitoxin system in *E. coli* during oxidative stress. . Genes with the highest and lowest expression in the single-cell transcriptomic analysis are indicated after treating exponentially-growing cells were with 20 mM H_2_O_2_ for 10 min. WT is BW25113. Largest values indicated by bold text.

| Gene | Cluster | WT | $\Delta mqsRA\Delta kan$ | Gene | Cluster | WT | $\Delta mqsRA\Delta kan$ |
| --- | --- | --- | --- | --- | --- | --- | --- |
| *yneL* | 1 | -0.7 | -0.9 | *yciF* | 1 | -0.26 | -0.83 |
|  | 2 | -4.4 | 1.5 |  | 2 | -5.12 | 2.15 |
|  | 3 | 2.3 | 1.9 |  | 3 | **8.76** | 0.23 |
|  | 4 | 2.93 | 0.28 |  | 4 | 3.39 | 0.39 |
|  | 5 | 3.08 | 0.43 |  | 5 | 3.54 | 0.54 |
|  | 6 | **9.21** | 0.53 |  | 6 | 3.89 | 0.64 |
|  | 7 | 3.54 |  |  | 7 | 4 |  |
| *gatR* | 1 | -0.07 | 1.82 | *yoeA* | 1 | 1.15 | -1.57 |
|  | 2 | -4.93 | -0.51 |  | 2 | -4.93 | 2.04 |
|  | 3 | 2.95 | 0.93 |  | 3 | 2.95 | 1.12 |
|  | 4 | **9.2** | 1.09 |  | 4 | 3.59 | 1.28 |
|  | 5 | 3.73 | 1.24 |  | 5 | 3.73 | 1.43 |
|  | 6 | 4.08 | 1.34 |  | 6 | 4.08 | 2.75 |
|  | 7 | 4.2 |  |  | 7 | **8.59** |  |
| *lfgA* | 1 | 0.74 | -0.57 | *ydiL* | 1 | 0.74 | 0.38 |
|  | 2 | -4.12 | 1.88 |  | 2 | -4.12 | 0.56 |
|  | 3 | 3.76 | 1.93 |  | 3 | 3.76 | 0.02 |
|  | 4 | 4.39 | 2.09 |  | 4 | 4.39 | 0.19 |
|  | 5 | 4.54 | 2.24 |  | 5 | **8.54** | 0.33 |
|  | 6 | 4.89 | 2.34 |  | 6 | 4.89 | 1.53 |
|  | 7 | **9** |  |  | 7 | 5 |  |
| *yagA* | 1 | 0.15 | -0.72 | *ibpA* | 1 | 0.162 | -0.068 |
|  | 2 | -4.71 | 1.3 |  | 2 | -0.166 | -0.404 |
|  | 3 | 3.18 | 1.12 |  | 3 | 0.008 | 1.272 |
|  | 4 | **8.98** | 1.28 |  | 4 | 0.16 | 1.218 |
|  | 5 | 4.22 | 2.65 |  | 5 | 0.196 | 0.106 |
|  | 6 | 4.57 | 1.53 |  | 6 | 0.11 | -0.334 |
|  | 7 | 4.68 |  |  | 7 | -0.092 |  |
| *holE* | 1 | -0.43 | -2.31 | *ibpB* | 1 | 0.632 | -0.55 |
|  | 2 | -4.12 | 3.62 |  | 2 | -0.666 | 1.034 |
|  | 3 | 2.59 | 1.6 |  | 3 | 0.398 | 0.974 |
|  | 4 | 3.22 | 1.77 |  | 4 | 0.574 | 0.862 |
|  | 5 | 3.37 | 1.92 |  | 5 | 0.466 | 0.898 |
|  | 6 | **8.89** | 2.02 |  | 6 | 0.726 | 0.896 |
|  | 7 | 3.83 |  |  | 7 | 0.322 |  |

Based on the single-cell transcriptome results, we tested 10 knockouts of the most highly-induced genes and found the *lfgA* deletion nearly completely prevented cells from surviving 20 mM H_2_O_2_ for 10 min (99.990 ± 0.004% death) whereas the wild-type strain had only 14 ± 13% death. Since we were unable to complement this phenotype by producing LfgA in a *lfgA* deletion mutant, we investigated whether a polar mutation was involved via kanamycin insertion into *lfgA* by investigating the next gene downstream of *lfgA*, *lfgB* (**Figure S1** and **Table S1**), and found deletion of *lfgB* also prevents survival with 20 mM H_2_O_2_ for 10 min (91 ± 8% death); this phenotype could be completely complemented by producing LfgB from pCA24N-*lfgB* (**Table 2**). Moreover, since RpoS positively controls KatG/KatE catalase activity^51^, these results were confirmed by observing the oxygen bubbles produced from catalase activity after incubating the *lfgB* mutant and complemented strain for 10 minutes with 20 mM of H_2_O_2_ (**Supplementary Figure S2**); quantifying the catalase results, the *lfgB* mutation reduced catalase activity by 62 ± 12% and producing LfgB from pCA24N-*lfgB* nearly completely restored catalase activity (94 ± 9%, **Fig. S2)**. Hence, we focused on LfgB to determine its role with MqsR/MqsA.

**Table 2.** Phenotypes of BW25113 (WT) and BW25113 Δ*lfgB* under different stresses. STD is standard deviation.

| Condition | Strain | % death | STD | Ratio |
| --- | --- | --- | --- | --- |
| **H2O2** | WT | 14 | 10 | 1 |
| | $\Delta lfgB$ | 91 | 8 | 6.4 |
| | $\Delta lfgB/pCA24N$ | 64 | 20 | 4.6 |
| | $\Delta lfgB/pCA24N-lfgB$ | 26 | 20 | 1.8 |
| **Acid** | WT | 28 | 4 | 1 |
| | $\Delta lfgB$ | 39 | 5 | 1.4 |
| | $\Delta lfgB/pCA24N$ | 35 | 4 | 1.3 |
| | $\Delta lfgB/pCA24N-lfgB$ | 25 | 2 | 0.9 |
| **Heat** | WT | -13 | 7 | 1 |
| | $\Delta lfgB$ | 18 | 11 | -1.4 |
| | $\Delta lfgB/pCA24N$ | 21 | 3 | -1.6 |
| | $\Delta lfgB/pCA24N-lfgB$ | -14 | 3 | 1.0 |

Since RpoS also controls the heat^49^ and acid response^2^ in *E. coli*, we hypothesized that inactivating LfgB should reduce viability after heat and acid treatments. Consistent with the reduction in the oxidative stress response, we found that the *lfgB* deletion reduces survival during acid stress (pH 4.0 for 10 min) stress (39 ± 5% death for *lfgB* vs. wild-type 28 ± 4% death for wild-type), as well as during heat (30 minutes at 50°C) stress (18% ± 12% death for *lfgB* vs. 0 ± 7% death for wild-type). Both phenotypes were complemented by producing LfgB from pCA24N-*lfgB* (**Table 2**).

Note that LfgB does not play a role in persister cell formation since survival after three hours with ampicillin at 10x the minimum inhibitory concentration there was little difference in cell viability (0.8 ± 0.4% viable for wild-type vs. 0.4 ± 0.2% viable for *lfgB*). Hence, LfgB is important for the stress response rather than antibiotic persistence. Also, deleting *lfgB* reduces the growth rate in LB medium by 25% (1.2 ± 0.1/h vs. 1.6 ± 0.2/h), so the dramatic reduction in viability of the *lfgB* mutant in the presence of H_2_O_2_ is not a result of poor growth.

LfgB is a poorly-characterized protein of cryptic prophage CP4-57 whose production previously led to a mutator phenotype LfgB^61^. To understand the relationship of this protein with the MqsR/MqsA TA system, we analyzed the RNA structure of the operon, finding two possible 5’-GCU sites accessible to toxin MqsR for *lfgB*^8^ in the predicted minimum free energy (MFE) structure for whole operon mRNA (**Figure S3A)**, which are not available in the MFE predicted structure of only the transcript containing just *lfgB* mRNA (**Figure S3B**). Hence, MqsR may degrade the mRNA containing *lfgB*.

### MqsA binds the *lfgA* promoter

We also considered the possibility that MqsA regulates the *lfg* operon by binding at its palindromic sequence 5’-ACCT N(2,6) AGGT upstream of the promoter as shown previously for the *mqsRA, csgD,* and *rpoS* promoters^6, 45, 51^. We found a probable MqsA palindromic sequence, 5’- ACCG (N5) CGGT, (grey highlight) 162 bp upstream of the start codon of *lfgA* (**Figure S4**). Thus, we hypothesized that MqsA represses transcription of the operon and overproduced MqsA from pCA24N- *mqsA* and observed that *lfgA* and *lfgB* are repressed 4 ± 1 and 3 ± 0.6-fold, respectively (**Table S2**). However, using EMSA, we found that mutating the *lfgA* promoter to interrupt this MqsA palindrome did not affect MqsA binding (**Fig. S5AB**). Hence, we conducted a DNA footprinting assay (**Fig. S6**) and determined the MqsA binding site is 245 bp upstream of the start codon, with a putative palindromic sequence 5’-ACAT (N2) ACAT (green highlight) (**Fig. S4**). Inactivating this MqsA-binding site in the *lfgA* promoter region via mutation confirmed the DNA footprinting results in that MqsA binding was abolished (**Fig. S5C**). Hence, MqsA, a known regulator, binds the promoter of the operon containing *lfgB*.

### LfgB controls the H_2_O_2_ response through MqsA degradation

To gain further insights into how LfgB interacts with the MqsR/MqsA TA system, we analyzed the protein structure of *lfgB*. Critically, LfgB is a putative zinc protease based on its predicted structure (UniProtKB - P52140), with a MPN domain (residues 38 – 160) and a JAMM motif (metalloprotease like zinc site)^3, 9^ (**Figure 1**). Based on this predicted structure, we purified LfgB and tested its protease activity against purified MqsA and found that LfgB degrades MqsA after overnight incubation at 37 °C (**Figure 2**). Unfortunately, the solubility of LfgB is extremely low, and we were unable to improve its solubility after many attempts, including purification under denaturing conditions and fusing SUMO and GST tags to LfgB.

**Fig. 1.**
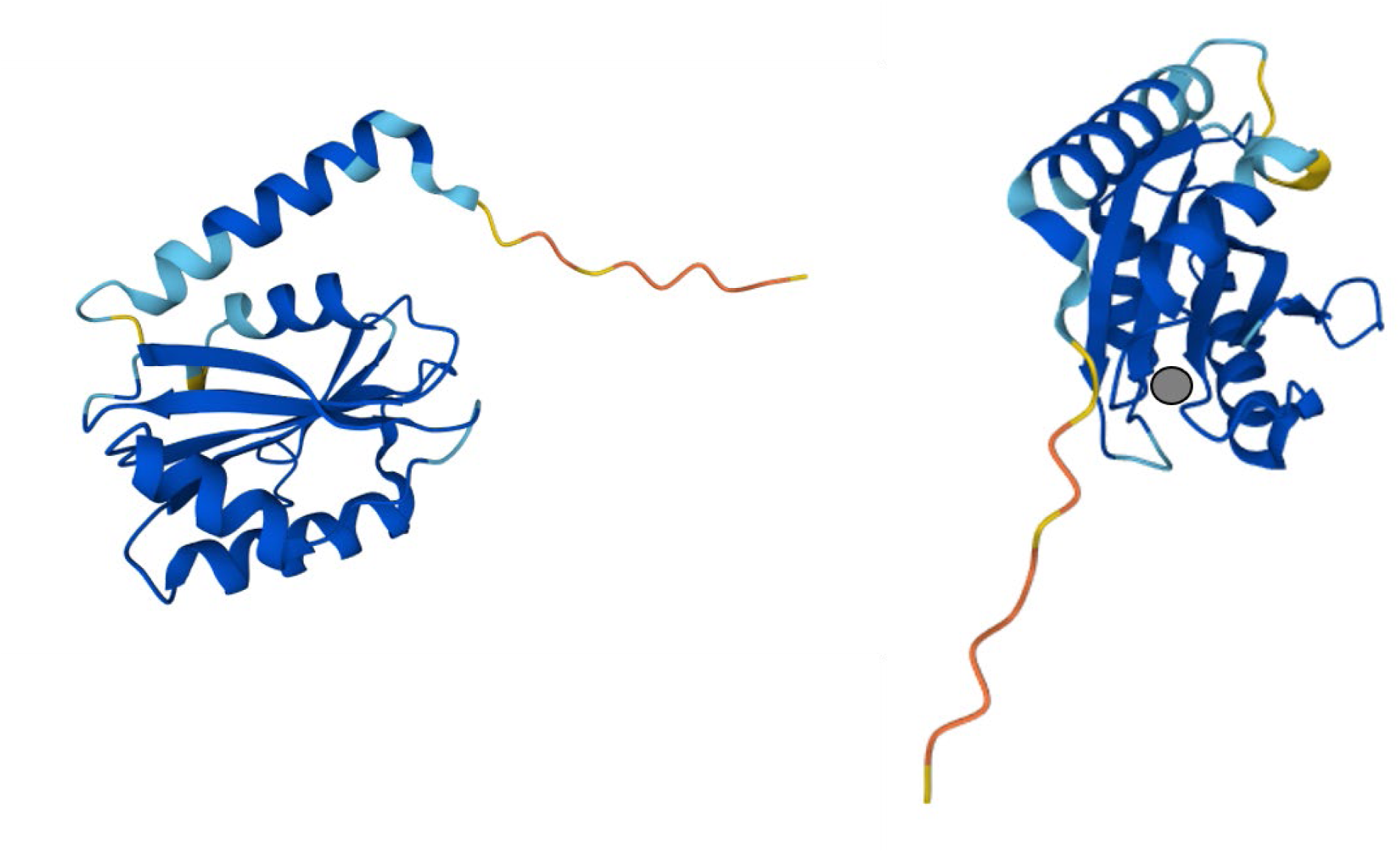
Two views of the predicted LfgB structure (UniProtKB - P52140). The grey circle represents zinc-binding residues His 109, His 111, and Asp 122.

**Fig. 2.**
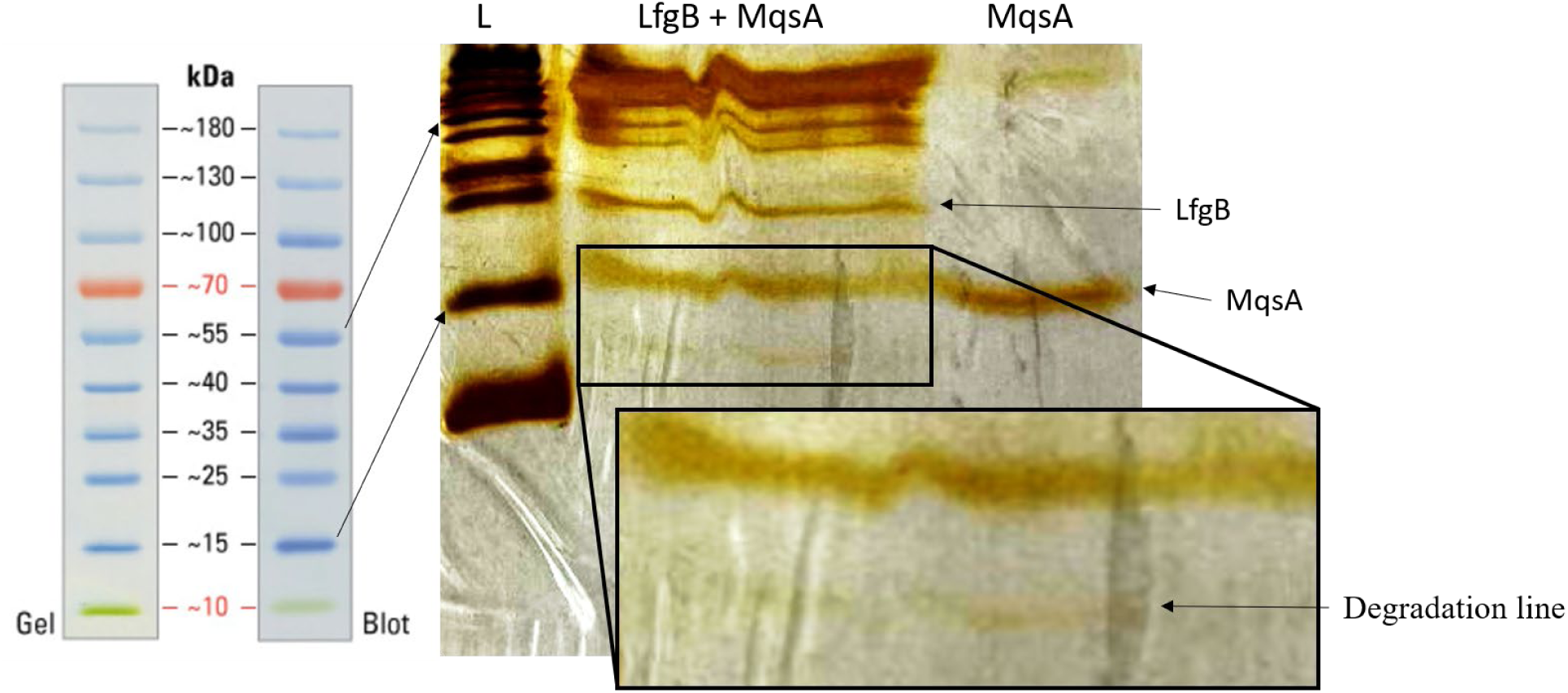
SDS-PAGE demonstrating degradation of MqsA by LfgB. Purified MqsA and LfgB were mixed in enzyme reaction buffer and incubated overnight at 37°C. L indicates ladder.

## DISCUSSION

Here, using the single cell transcriptome for the first time to study TA systems, we determined additional insights into how the MqsR/MqsA type II TA system is physiologically important for the growth of *E. coli* during exposure to H_2_O_2_ stress. Specifically, we (i) identified the *lfg* operon of cryptic prophage CP4-57 is induced during oxidative stress in single cells, (ii) found MqsA represses the *lfg* operon, and (iii) characterized LfgB as a protease that degrades antitoxin MqsA. Remarkably, our results demonstrate that the cell combines the tools of its former enemy, prophage CP4-57, with that of the MqsR/MqsA TA system, to regulate its stress response.

Cryptic prophage CP4-57 has been linked to *E. coli* cell growth, biofilm formation, motility, and carbohydrate metabolism^53^, and we found previously that the *lfgB* and *lfgA* deletions increase biofilm formation 6-fold and 2-fold, respectively^53^. In addition, we found the *lfgD* mutation reduces MqsR toxicity^15^. Therefore, by characterizing protease LfgB, our results provide additional proof that cryptic prophages are beneficial and are involved in the stress response^52^.

Our results also increase add another facet to MqsA regulation by finding a new protease that degrades MqsA. As previously demonstrated, Lon protease can degrade MqsA as well as other antitoxins under oxidative stress^51^. In addition, ClpXP degrades MqsA in the absence of the zinc that is used to stabilize the structure of MqsA; i.e., when it is unfolded^50^. It was proposed that the ClpX recognition site is accessible under non-stress conditions; however, under oxidative conditions, cysteine residues are oxidized preventing the correct folding and the binding of zinc, allowing ClpXP to degrade MqsA^50^. Hence, our results with protease LfgB provide additional evidence for the selective degradation of free antitoxins under stress conditions^30, 50, 51^.

Our proposed mechanism is shown in **Figure 3**. In the absence of stress, one physiological role of MqsA is to inhibit *rpoS* transcription^51^, which is important for rapid growth. However, under stress conditions (H_2_O_2_, acid, heat), Lon protease^51^, ClpXP protease^50^, and LfgB protease degrade antitoxin MqsA, facilitating the formation of RpoS and activation of the stress response. This also shifts the balance to MqsR^39, 51^, which then performs differential mRNA decay^23^, based on the presence of single-stranded, 5’-GCU sites^8^. One example of the differential mRNA decay is the degradation of the transcript for antitoxin GhoS, which results in activation of toxin GhoT (whose transcript lacks 5’-GCU sites)^54^; this then allows toxin GhoT to reduce ATP and growth^7^.

**Fig. 3.**
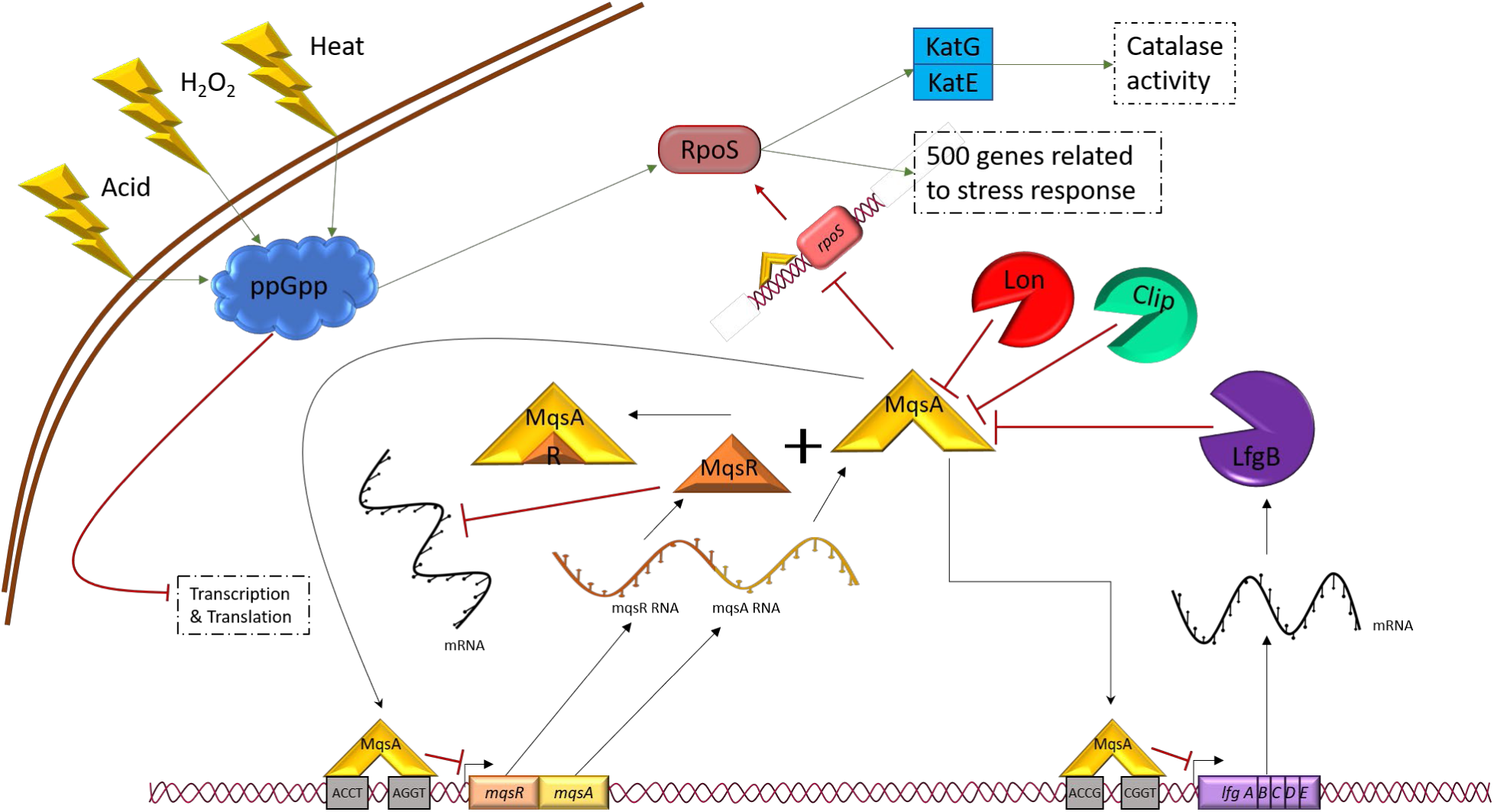
Scheme for the MqsR/MqsA toxin/antitoxin/LfgB protease stress response mechanism, and its relationship with MqsA. Green arrows indicate activation, and red lines indicate inhibition.

Therefore, the type II TA system MqsR/MqsA is a multi-faceted regulator that facilitates growth of *E. coli* populations residing in the gut during exposure to bile (oxidative) stress. Since bile plays an important role as an interkingdom signal in the GI tract^21^, our results also illustrate how a TA system can play an important role in host-microbe interactions by ensuring the survival of a commensal bacterium.

## EXPERIMENTAL PROCEDURES

### Bacterial strains and growth conditions

The *E. coli* K-12 strains and plasmids used in this study are listed in **Table 3**. All cultures were grown in lysogeny broth (LB) medium^41^ at 37°C with 30 μg/ml of chloramphenicol to maintain the pCA24N plasmids.

**Table 3.** Bacterial strains and plasmids used in this study. Km^R^ indicates kanamycin resistance and Cm^R^ indicates chloramphenicol resistance.

| Strain | Genotype | Source |
| --- | --- | --- |
| BW25113 | *rrnB3 ΔlacZ4787 hsdR514 Δ(araBAD)567 Δ(rhaBAD)568 rph-1* | 1 |
| BW25113 *ΔmqsRA Δkan* | BW25113 *ΔmqsRA* ΔKm<sup>R</sup> | 23 |
| BW25113 *ΔlfgA* | BW25113 *ΔlfgA* Ω Km<sup>R</sup> | 1 |
| BW25113 *ΔgatR* | BW25113 *ΔgatR* Ω Km<sup>R</sup> | 1 |
| BW25113 *ΔyneL* | BW25113 *ΔyneL* Ω Km<sup>R</sup> | 1 |
| BW25113 *ΔynfP* | BW25113 *ΔynfP* Ω Km<sup>R</sup> | 1 |
| BW25113 *ΔyagA* | BW25113 *ΔyagA* Ω Km<sup>R</sup> | 1 |
| BW25113 *ΔholE* | BW25113 *ΔholE* Ω Km<sup>R</sup> | 1 |
| BW25113 *ΔyoeA* | BW25113 *ΔyoeA* Ω Km<sup>R</sup> | 1 |
| BW25113 *ΔyidL* | BW25113 *ΔyidL* Ω Km<sup>R</sup> | 1 |
| BW25113 *ΔibpA* | BW25113 *ΔibpA* Ω Km<sup>R</sup> | 1 |
| BW25113 *ΔibpB* | BW25113 *ΔibpB* Ω Km<sup>R</sup> | 1 |
| BW25113 *ΔlfgB* | BW25113 *ΔlfgB* Ω Km<sup>R</sup> | 1 |
| BW25113 *ΔypjJ* | BW25113 *ΔypjJ* Ω Km<sup>R</sup> | 1 |
| **Plasmid** |  |  |
| pCA24N | Cm<sup>R</sup>; *lacI*<sup>q</sup>, pCA24N | 25 |
| pCA24N-*lfgA* | Cm<sup>R</sup>; *lacI*<sup>q</sup>, pCA24N P<sub>T5-lac</sub>::*lfgA* | 25 |
| pCA24N-*lfgB* | Cm<sup>R</sup>; *lacI*<sup>q</sup>, pCA24N P<sub>T5-lac</sub>::*lfgB* | 25 |
| pCA24N-*mqsA* | Cm<sup>R</sup>; *lacI*<sup>q</sup>, pCA24N P<sub>T5-lac</sub>::*mqsA* | 25 |
| pET28b | Km<sup>R</sup>, expression vector with T7 promoter | Novagen |
| pET28b-*mqsA* | Km<sup>R</sup>, *lacI*<sup>q</sup>, pET28b P<sub>T7-lac</sub>::*mqsA* with *mqsA* C-terminus His-tagged | this study |

### Single-cell transcriptome analysis

BW25113 and its unmarked isogenic mutant Δ*mqsRA* were harvested during exponential growth (turbidity 0.8 nm at 600 nm), treated with 20 mM H2O2 for 10 min, fixed with formaldehyde (1%) for 30 min. After centrifugation, cell pellets were washed with PBS and resuspended in 4:1 vol% methanol:glacial acetic acid, and analyzed at the single-cell level as described previously^33^.

### Viability assays with hydrogen peroxide, acid, and heat

Cells were cultured in LB to a turbidity of 0.8 at 600 nm, then exposed to 20 mM H_2_O_2_ for 10 min, to acid conditions (pH 4) for 10 min, or to heat (50 °C) for 30 minutes. For cyclic exposure to acid (pH 2.5), cells were exposed in four times for 10 min/cycle with one hour growth in between each treatment.

### Persister cell formation^28^

Overnight cultures were grown to a turbidity of 0.8 at 600 nm, then cells were resuspended in LB-ampicillin (100 μg/mL, 10 MIC) and incubated for 3 h. Cells were washed twice with PBS, and viable cells were quantified using serial dilution and spot plating onto LB agar plates. Experiments were performed with at least three independent cultures.

### RNA structure prediction and DNA palindrome search

The RNA predicted structures and palindrome search were obtained using the NCBI *E. coli* BW25113 genome sequence (NZ_CP009273.1), and the MFE RNA structures were predicted by the RNAfold webserver (http://rna.tbi.univie.ac.at/cgi-bin/RNAWebSuite/RNAfold.cgi).

### qRT-PCR

Overnight cultures of Δ*mqsRA*/pCA24N and Δ*mqsRA/*pCA24N-*mqsA* were grown to a turbidity of 0.1 at 600 nm in LB/chloramphenicol medium, then 1 mM of IPTG for 30 min was used to induce expression of *mqsA*. Cells were rapidly cooled in ethanol/dry ice, centrifuged, and the pellets were collected with RNALater buffer (Applied Biosystems, Foster City, CA, USA), to stabilize RNA. The RNA was purified using the RNA purification kit (Roche). qRT-PCR were performed following the manufacturer’s instructions for the iTaq™ Universal SYBR® Green One-Step Kit (Bio-Rad) using 100 ng of total RNA as template. Primers were annealed at 60°C and data were normalized against the housekeeping gene *rrsG*^45^. The specificity of the qRT-PCR primers (**Table S3**) was verified via standard PCR, and fold changes were calculated using the method of Pfaffl^38^ using the 2^-ΔΔCT^.

### Proteolytic assay

Purified MqsA and LfgB were mixed in enzyme reaction buffer (40 mM HEPES-KOH, 25 mM Tris-HCl, 4% sucrose, 4 mM DTT, 11 mM magnesium acetate, and 4 mM ATP) and incubated overnight at 37°C. SDS-PAGE was conducted using 5% stacking and 18% acrylamide resolving sections and staining following the manufacturer’s instructions (Pierce Silver Stain kit, Thermo Scientific).

### MqsA purification and electrophoretic mobility shift assay (EMSA)

The *mqsA* coding region was amplified with primer pair pET28b-*mqsA*-F/R using MG1655 genomic DNA as the template. The amplified DNA fragment was purified, quantified, and ligated into pET28b digested with NcoI/HindIII. pET28b- *mqsA* was used to purify MqsA using standard methods^19^. For DNA probes to investigate MqsA binding, the promoter region of *yfjY* was amplified with primer pair *yfjY*-P-F and *yfjY*-P-R, and the two mutant probes were also amplified with primer pairs *yfjY*-MP-F/*yfjY*-P-F and *yfjY*-MP2-F/*yfjY*-P-F (**Table S3**). The probes were purified and labelled with biotin by using the Biotin 30-End DNA Labeling Kit (Thermo Scientific, Rockford, USA)f, and 0.25 pmol were used to assay the binding reaction with a series of concentrations of MqsA^31^. The stopped reaction mixtures were run on a 6% polyacrylamide gel in Tris-borate EDTA and were then transferred to nylon membranes. The Chemiluminescence Nucleic Acid Detection Module Kit (Thermo Scientific) was used to observe the shift of the DNA probes on the membranes.

### DNase I footprinting assay

This assay was conducted as reported previously^31^.The FAM-labelled probe covering the promoter region of *yfjY* was amplified with primer pair FAM-*yfjY*-P-F and *yfjY*-P-R, and the products was purified with QIAEX II Gel Extraction Kit (Qiagen, Hilden, Germany). The labelled probes (200 ng) were mixed with varying amounts of MqsA, and the mixtures were incubated for 30 min at 25°C. An orthogonal combination of DNase I (NEB, M0303S) and incubation time were used to achieve the best cutting efficiency. A final concentration of 200 mM EDTA was added to the reaction mixture to stop the reaction. The DNA was purified again with a QIAEX II Gel Extraction Kit (Qiagen, Hilden, Germany), and the generated products were screened and analyzed as reported^31^.

### Catalase assay

Catalase activity was determined spectrophotometrically by recording the decrease in the absorbance of H_2_O_2_ at 240 nm in a UV/VIS spectrophotometer as described previously^14^. Briefly, five independent cultures per strain were grown overnight, 1 mL aliquots were taken, cells were collected by centrifugation for 1 min at 13,000 rpm, washed with sterile HEPES buffer (50 mM, pH 7.5), centrifuged again, frozen with liquid nitrogen, and stored at -70 °C. Thawed pellets were resuspended in 1 mL of sterile cold HEPES buffer (50 mM, pH 7.5) with MgCl_2_ 10 mM and 0.025% triton X-100 and disrupted by sonication using 2 pulses of 20 sec with 1 minute pause between cycles. Catalase activity was determined using 15 mM H_2_O_2_ as substrate and normalized based on the protein level in the cell extracts as determined using the Bradford method.

## ACKNOWLEDGEMENTS

This work was supported by a Fulbright Scholar Fellowship for LFG. We are also grateful for the Keio and ASKA strains provided by the National Institute of Genetics of Japan and the single cell transcriptomic experiment by International Flavors and Fragrances.

## SUPPLEMENTARY INFORMATION

**Table S1.** Single cell expression levels for *mqsRA* vs. wild-type in the presence of 20 mM H_2_O_2_ stress for the *lfg* operon (*lfgA* is also in Table 1).

| Gene | Cluster | WT | $\Delta mqsRA \Delta kan$ |
| --- | --- | --- | --- |
| *lfgA* | 1 | 1.5 | -0.8 |
|  | 2 | -2.0 | 1.9 |
|  | 3 | 3.7 | 1.8 |
|  | 4 | 4.3 | 1.7 |
|  | 5 | 4.9 | 1.9 |
|  | 6 | 4.8 | 2.0 |
|  | 7 | 5.7 |  |
| *lfgB* | 1 | 1.1 | 0.5 |
|  | 2 | -0.5 | 0.01 |
|  | 3 | 4.1 | 2.6 |
|  | 4 | 4.8 | 2.8 |
|  | 5 | 4.9 | 2.9 |
|  | 6 | 5.3 | 3.8 |
|  | 7 | 5.4 |  |
| *lfgC* | 1 | 3.2 | -1.3 |
|  | 2 | -2.6 | 1.8 |
|  | 3 | 3.8 | 1.9 |
|  | 4 | 4.4 | 2.9 |
|  | 5 | 4.6 | 2.2 |
|  | 6 | 4.9 | 2.3 |
|  | 7 | 5.0 |  |
| *lfgD* | 1 | 1.8 | 1.3 |
|  | 2 | -1.9 | -0.2 |
|  | 3 | 3.1 | 1.2 |
|  | 4 | 4.0 | 1.6 |
|  | 5 | 4.2 | 1.7 |
|  | 6 | 4.3 | 1.6 |
|  | 7 | 4.8 |  |
| *lfgE* | 1 | 1.1 | -0.8 |
|  | 2 | -1.2 | 1.0 |
|  | 3 | 2.5 | 1.1 |
|  | 4 | 3.1 | 1.4 |
|  | 5 | 3.8 | 1.3 |
|  | 6 | 3.8 | 1.9 |
|  | 7 | 3.6 |  |

**Table S2.** Repression of *lfgAB* in the BW25113 *mqsRA* deletion mutant during production of MqsA. MqsA was produced by 1 mM IPTG for 30 min.

| Gene | *rrsG* |  | *lfgA* |  | *lfgB* |  |
| --- | --- | --- | --- | --- | --- | --- |
| Strain | $\Delta mqsRA$<br>pCA24N | $\Delta mqsRA$<br>pCA24N-*mqsA* | $\Delta mqsRA$<br>pCA24N | $\Delta mqsRA$<br>pCA24N-*mqsA* | $\Delta mqsRA$<br>pCA24N | $\Delta mqsRA$<br>pCA24N-*mqsA* |
| CT Mean | 13.1 ± 0.6 | 11.6 ± 0.4 | 22.7 ± 0.5 | 23 ± 1 | 24.7 ± 0.9 | 24.7 ± 0.4 |
| $\Delta Ct$ | | | 9.6 | 11.7 | 11.6 | 13.1 |
| $\Delta\Delta Ct$ | | | | 2 ± 1 | | 1.5 ± 0.6 |
| Fold change |  |  |  | -4 |  | -2.8 |

**Table S3.**
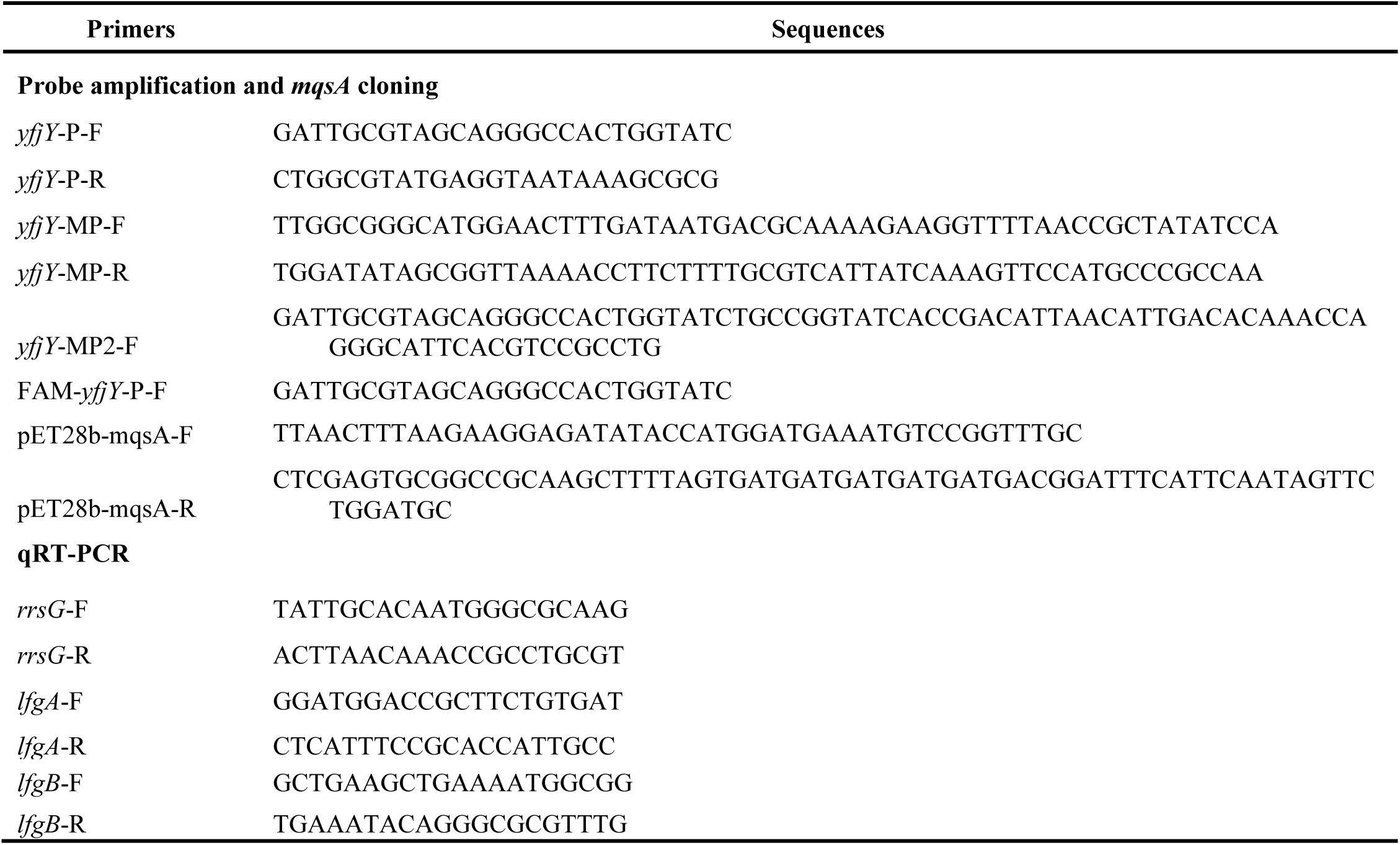
Primers used in this study. F indicates forward primer and R indicates reverse primer. P indicates promoter, MP indicates mutant promoter.

**Figure S1.**
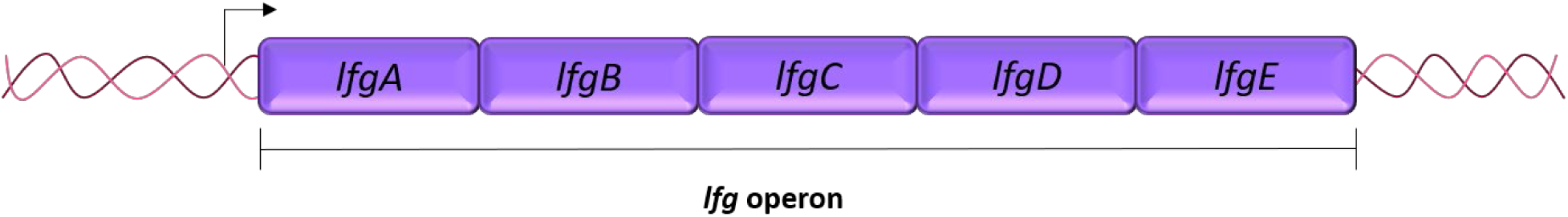
Scheme representing the organization of the *lfg* operon inside cryptic prophage CP4-57.

**Figure S2.**
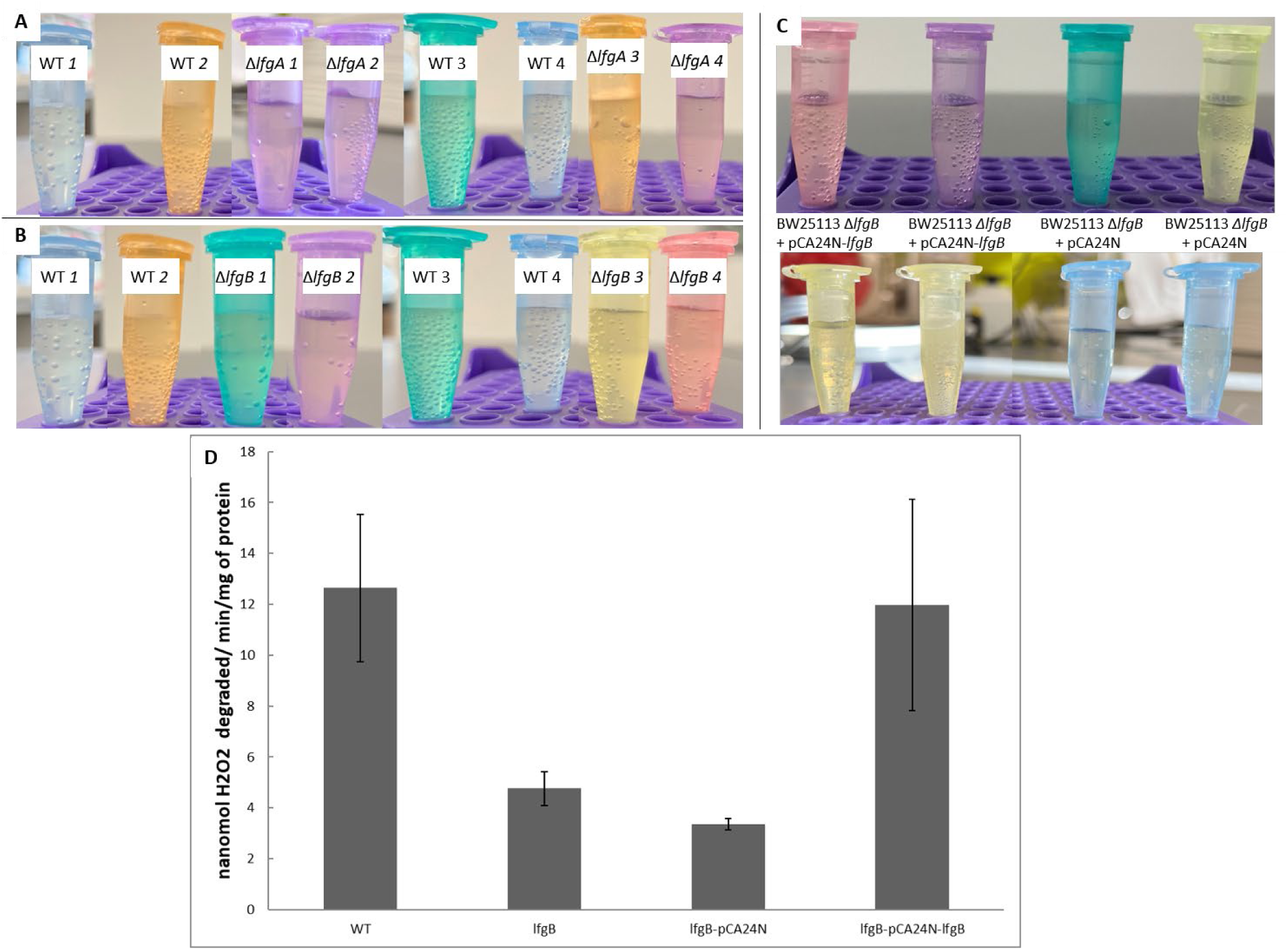
Images of oxygen bubble formation from catalase activity during the hydrogen peroxide assay for (**A**) BW25113 WT compared to BW25113 Δ*lfgA*, (**B**) BW25113 WT compared to BW25113 Δ*lfgB*, (**C**) BW25113 pCA24N compared to BW25113 pCA24N-*lfgB*, (**D**) quantified catalase activity.

**Figure S3.**
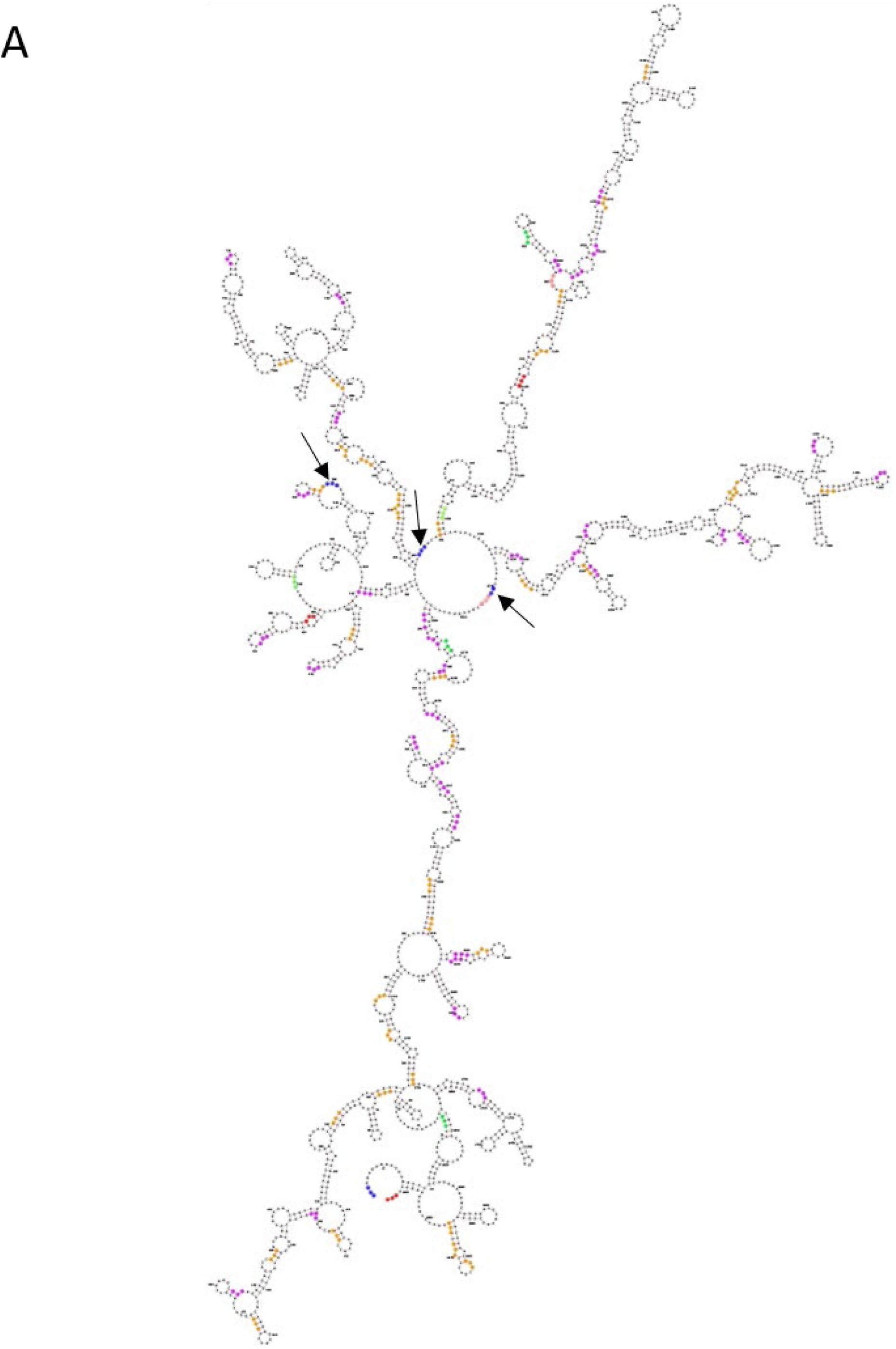
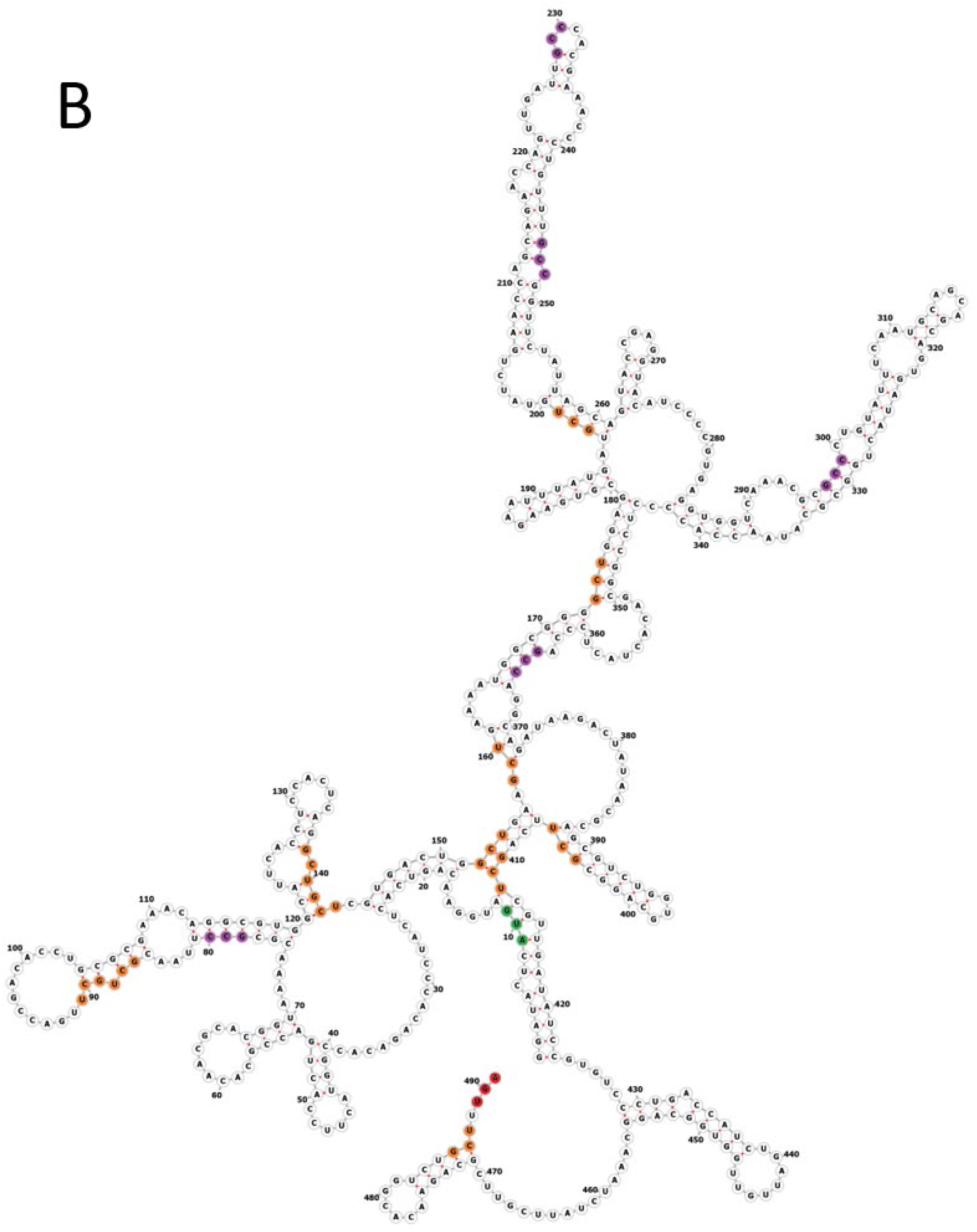
Predicted (MFE) RNA secondary structures for (**A**) the *lfg* operon and (**B**) *lfgB* indicating the presence of MqsR cleavage sites (5’-GCU). Green circles indicate start codons, red and pink circles indicate the termination codons, blue circles indicate 5’-GCU MqsR cleavage sites inside a stem-loop, purple circles indicate 5’-GCU sites outside stem-loops, and orange circles indicate 5’-GCC sites. Arrows indicate likely MqsR single-stranded sites.

**Figure S4.**
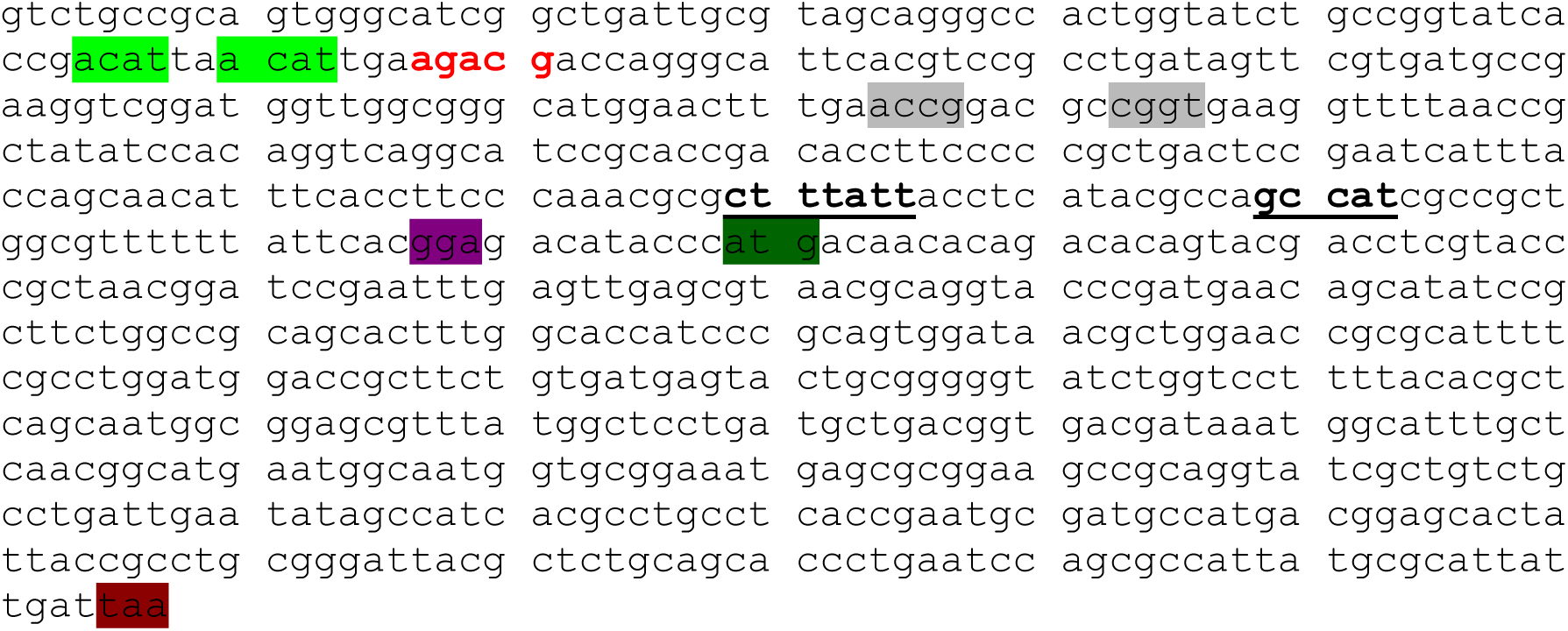
Sequence of the first gene of the *lfg* operon (*lfgA*) along with the upstream region (200 bp). MqsA-binding region identified from DNA footprinting (bold red), and previously described as 5’-ACCT N(2,6) AGGT^51^ (highlighted in gray). Bold and underlined indicates the -35 and -10 promoter regions, purple highlight indicates the ribosome binding site, and green highlight indicates the *lfgA* start codon. The actual binding palindrome is highlighted in green.

**Fig. S5.**
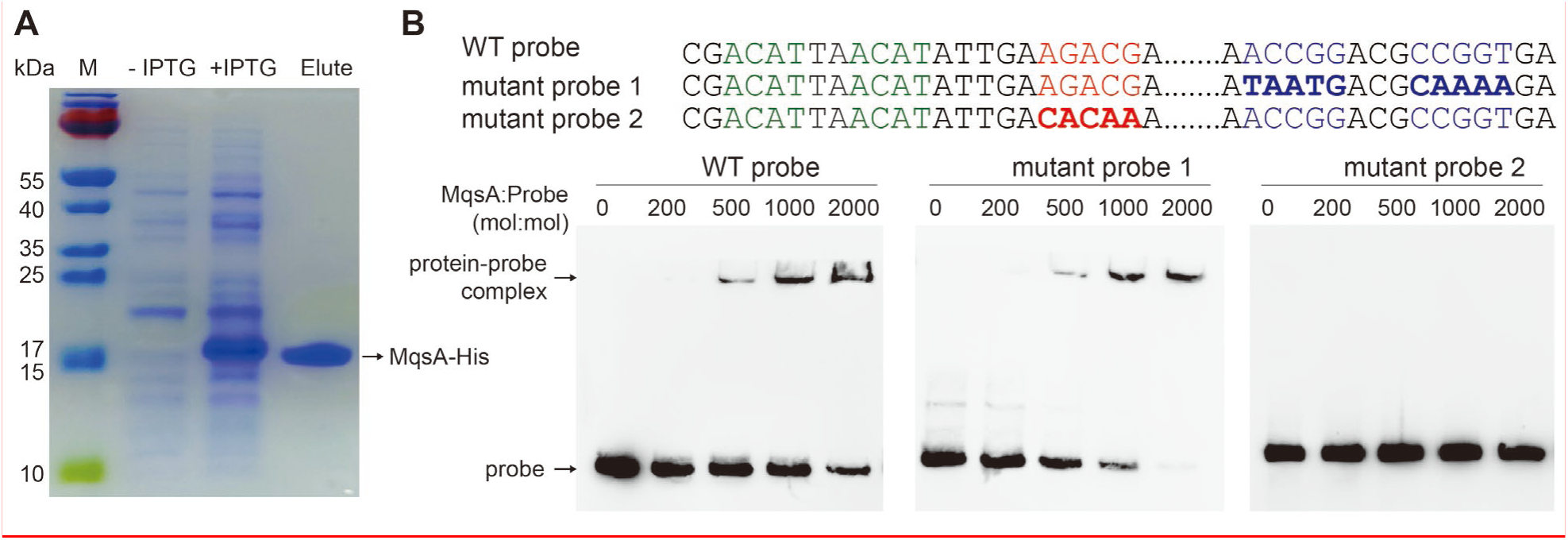
**EMSA shows DNA-binding regulator MqsA binds the *lfg* operon**. **(A)** Purification of MqsA. **(B)** Mutating the *lfgA* promoter to interrupt the MqsA palindrome 5’-ACCG (N5) CGGT did not affect MqsA binding (left and middle, mutant probe 1), but mutating the region identified by the DNA footprinting assay (bold red in **Fig. S6**) 252 bp upstream of the start codon, with nearby putative palindromic sequence 5’-ACAT (N2) ACAT (green highlight in **Fig. S6**), abolishes MqsA binding (right).

**Fig. S6.**
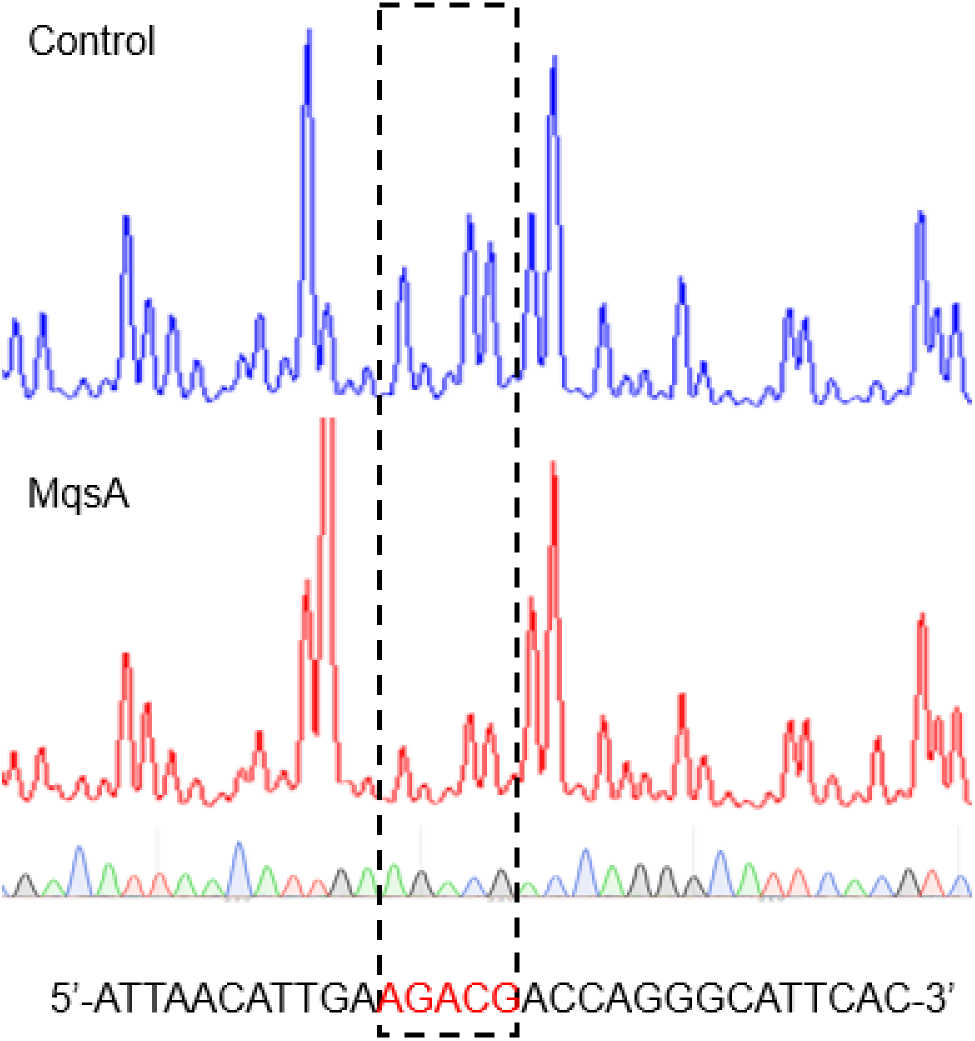
DNA footprinting. DNA footprinting shows the MqsA binding site is 252 bp upstream of the start codon of the *lfg* operon (bold red in **Fig. S4**), near the palindromic sequence 5’-ACAT (N2) ACAT (green highlight in **Fig. S4**).

